# Dung beetles do not profit from enhanced spatial heterogeneity in production forests: a large-scale forest manipulation experiment

**DOI:** 10.1101/2025.08.18.670815

**Authors:** Johanna Asch, Michael Scherer-Lorenzen, Kerstin Pierick, Clara Wild, Julia Rothacher, Jörg Müller, Orsi Decker, Simone Cesarz, Nico Daume, Jörn Buse, Marcell K. Peters

**Affiliations:** Department of Animal Ecology and Tropical Biology, Biocenter, University of Würzburg, Würzburg, Germany; Faculty of Biology, University of Freiburg, Geobotany, D-79104 Freiburg, Germany; Spatial Structures and Digitization of Forests/Silviculture and Forest Ecology of the Temperate Zones, University of Göttingen, Göttingen, Germany; Chair of Conservation Biology and Forest Ecology, Biocenter, University of Würzburg, Rauhenebrach, Germany; Bavarian Forest National Park, Grafenau, Germany; German Centre for Integrative Biodiversity Research (iDiv) Halle-Jena-Leipzig, Leipzig, Germany; Institute of Biology, Leipzig University, Leipzig, Germany; Berchtesgaden National Park, Berchtesgaden, Germany; Black Forest National Park, Seebach, Germany; Animal Ecology Group, BIOM, University of Bremen, Bremen, Germany

**Keywords:** β-diversity, dung beetles, climate, decomposition, forest management, forest experiment, spatial heterogeneity

## Abstract

1. Central European forest management strategies promoting structurally homogeneous closed-canopy forests have led to landscape-level declines in biodiversity and ecosystem multifunctionality. Conservation-targeted management programs aim at reintroducing structural heterogeneity into production forests, however, it is not well understood if species diversity and ecosystem functions generally profit from management promoting heterogeneity in forest structure. By removing and processing mammalian dung remains, dung beetles play an integral role in ecosystem functions in forests.
2. In one of the largest manipulative forest experiments in Central Europe to date, we analysed the effects of enhanced structural heterogeneity in production forests, climate, and mammalian defecation on dung beetle diversity and dung removal rates at the local and landscape level. We assessed communities of dung beetles and dung removal rates on 234 study patches (50 x 50 m) in eleven paired forest landscapes across a climatic gradient in Germany. Forest landscapes were either managed to conserve a homogeneous closed canopy or to create a heterogeneous forest structure with forest patches varying in canopy coverage and dead wood availability.
3. We did not find that more heterogeneously managed forests had higher dung beetle species diversity and dung removal rates. Canopy openings did not increase species turnover but decreased species diversity.
4. Along the climate gradient, dung beetle average biomass and dung removal decreased with increasing temperature. Canopy openings in combination with higher temperatures negatively impacted all abundant dung beetle species, but especially the large species *Anoplotrupes stercorosus*, comprising > 90% of the total dung beetle biomass.
5. *Synthesis and application:* Our results suggest that dung beetles do not profit from management increasing the structural heterogeneity of forests. Due to a restricted climate and habitat niche of the current Central European dung beetle fauna, future warming and openings in forest structure might have negative effects on dung beetles and consequently on the ecosystem services they provide in Central European production forests.

## Introduction

With the intensification of timber production and the introduction of continuous cover forestry, modern forestry has introduced profound changes to temperate production forests, shaping forest structure towards closed-canopy forests with little structural heterogeneity (Gossner et al., 2013; Pommerening & Murphy, 2004). As a result this homogenization of forests has contributed to biodiversity loss and, consequently, a loss of ecosystem services (Schelhaas et al., 2003; Seibold et al., 2019).

Novel experimental forest management concepts, such as the enhancement of structural beta complexity (ESBC) approach, aim at emulating the heterogeneous forest structure of natural forests. This can be achieved by varying canopy density and increasing the amount of different types of dead wood through silvicultural manipulations (Asbeck et al., 2023; Müller et al., 2023). Canopy openings and dead wood enrichments often have positive effects on local-scale (α-) biodiversity (Doerfler et al., 2018; Dove & Keeton, 2015; Lettenmaier et al., 2022; Rothacher et al., 2025). However, how structural heterogeneity influences landscape-scale biodiversity (γ-diversity), species turnover between local patches (β-diversity), and ecosystem functioning in forest landscapes is still little understood (Müller et al., 2023; Uhl et al., 2024).

Coprophagous beetles (dung beetles) are useful ecological indicators for changing environmental conditions as their community structure is strongly controlled by habitat and climate (Nichols & Gardner, 2011). Dung beetle communities in temperate forests and grasslands, for example, have been found distinct with little overlap of species between the two habitats (Buse & Entling, 2020; Frank et al., 2017). Dung beetles provide a variety of key ecosystem functions through dung removal, such as nutrient cycling, bioturbation, secondary seed dispersal, and parasite suppression (Nichols et al., 2008). This makes them particularly suited for studying the effects of forest structural complexity on species communities important for ecosystem functioning.

Because of their economic value in agricultural pasture systems, dung beetle communities in Europe have mostly been studied in grasslands. Although several studies have shown how land-use change has negatively affected coprophagous beetles on pastures and arable land, e.g., through the reduction of grazing livestock (Englmeier et al., 2022; Rosenlew & Roslin, 2008; Sánchez-Bayo & Wyckhuys, 2019), little is known about the effects of forest management on temperate forest communities (but see Frank et al. (2017) and Staab et al. (2022)).

The rise of global air temperatures exposes coprophagous beetles to climatic changes in combination with changing environments through land use. Warming climate could positively impact dung beetles as studies show that dung beetle abundance and α-diversity increase from colder to warmer environments in temperate forests (Staab et al., 2022; von Hoermann et al., 2022). In laboratory experiments, dung removal by large temperate dung beetle species increased when they were exposed to climate warming scenarios (Nervo et al., 2024). Given forest canopies have a temperature buffering effect, forest species are less exposed to changes in ambient temperature than open land communities (De Frenne et al., 2019). However, the increase in disturbance events may change forest structure towards more openings of the forest canopies (Seidl et al., 2017). Therefore, studying the effects of changes in forest structure in interaction with climatic conditions on forest communities is not only important for testing forest management strategies but also for predicting potential responses to future changes.

Since dung beetles mainly rely on mammalian dung for brooding, the abundance of large mammals is assumed to influence dung beetle species richness and abundance by controlling resource availability (Raine & Slade, 2019). Structural complexity has been shown to positively influence functional diversity of mammals and ungulates have been observed to prefer foraging in canopy gaps (Kuijper et al., 2009; Sukma et al., 2019). Therefore, forest structural heterogeneity might also have indirect effects on dung beetle communities by impacting mammal defecation rates.

Here we aim to disentangle the effects of forest management promoting structural heterogeneity, climate, and mammalian activity on dung beetle communities in temperate production forests in Central Europe. By assessing dung beetle communities and dung removal in 234 forest patches (50 x 50 m) in eleven distinct forest pairs in Germany, we compared conventional closed-canopy production forests to forests in which we enhanced the structural complexity by experimentally altering canopy openness and dead wood availability (ESBC-management). Additionally, we correlated mammal occurrences with dung beetle community composition and dung removal. We hypothesized that:

1. ESBC-management fosters higher β-diversity among forest patches and, in turn, leads to higher γ-diversity of dung beetles compared to conventionally managed control forests at the landscape scale.
2. By fostering higher β-diversity, ESBC-management leads to stronger differences in dung removal between forest patches and overall, to higher dung removal compared to conventionally managed control forests.
3. Dung removal is controlled by dung beetle α-diversity and biomass, and both dung removal and dung beetle α-diversity will be maximized in forests with warmer climates, as well as in habitats with high mammalian dung availability.

## Methods

### Study area and study design

This study was conducted within the research unit BETA-FOR (Müller et al. 2022) on 234 study patches distributed over eleven experimental forest landscapes (forest sites) in six regions across Germany. The forest landscapes spanned an elevational gradient from 38 m (Luebeck) to 1143 m (Bavarian Forest National Park) above sea level (Figure 1a). All forest sites were production forests of native broad-leaved trees with a dominance of European beech (*Fagus sylvatica*), European Ash (*Fraxinus excelsior*), and/or European hornbeam (*Carpinus betulus*). Each forest site was comprised of two differently managed forest districts composed of nine individual patches (50 x 50 m), or, in one region (University Forest), of 15 individual patches (Figure 1b, Figure S1). The patches in one forest district were all conventionally managed in the same way, i.e., with continuous cover forestry (thereafter, called control districts). Sites located in national parks resembled production forest sites. Patches in the paired district were subjected to one of a variety of experimental silvicultural manipulations (ESBC-treatments) for enhancing the structural complexity at the landscape scale (hereafter called ESBC districts) (Figure1b). ESBC-treatments differed in the way trees were manipulated: either in the center of the forest patch (aggregated ESBC-treatment), creating one continuous canopy opening, or tree manipulations were evenly distributed, reducing the density of the canopy, but not creating a continuous opening (distributed ESBC-treatment). ESBC districts were created in 2015 in the Bavarian Forest National Park (BF) and Passau (Pas), 2016 in the forests located in Saarland (Saa), Hunsrück (Hun), and Lübeck (Lue), and 2018 in the University Forest (UF). For an in-depth description of the BETA-FOR project and establishment of the forest sites, see Müller et al. (2022) and Müller et al. (2023).

**Figure 1.**
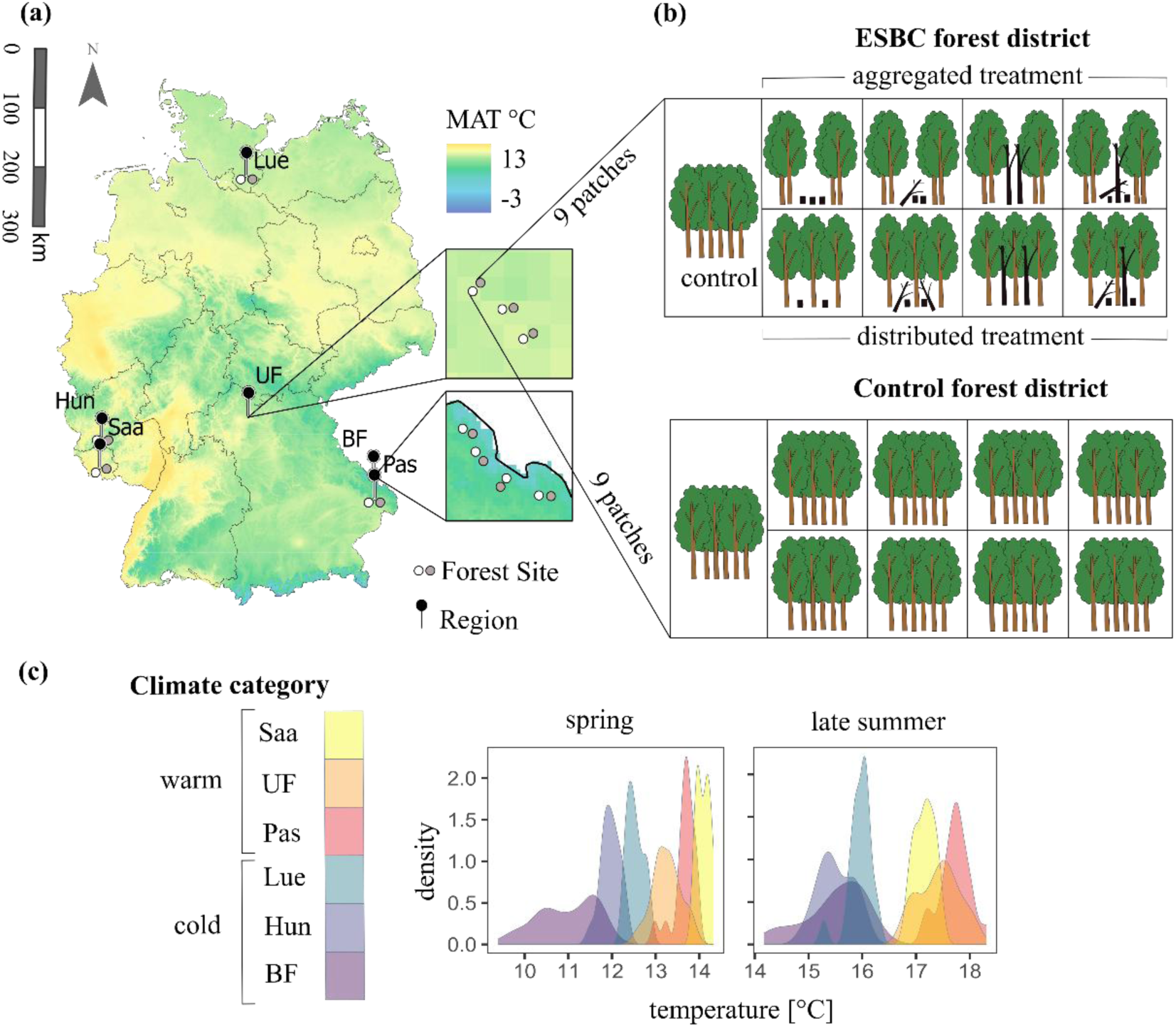
(a) Study regions and forest sites of the BETA-FOR research project, map colors represent 5-year averages of mean annual temperatures (2019 – 2023) (Source: Deutscher Wetter Dienst DWD). **(b)** Graphical representation of forest structure manipulations on each forest patch in the ESBC forest district (grey points on map) and control forest district (white points on map) at each forest site. Black color represents different dead wood manipulations. **(c)** Distributions of mean air temperatures during the sampling time of this study. Colors represent the different study regions, which were categorized into regions with cold or warm climate. Cold climate: BF, Bavarian Forest; Hun, Hunsrück; Lue, Lübeck; Warm climate: Pas, Passau; Saa, Saarland; UF, University Forest.

The forest regions were allocated to two climatic categories: warm climate (UF, Saa, Pas) and cold climate (Lue, Hun, BF), based on distributions of mean air temperatures during sampling (cold climate < 16.5 °C, warm climate > 16.5°C during summer sampling) (Figure 1c). Warmer regions additionally had lower average precipitation sums when comparing 5-year averages (2019 – 2023) (Figure S2).

### Decomposition rates

Decomposition rates were measured by exposing two pats of cow dung per plot for two days in spring (May) and late summer 2023 (mid-August – mid-September) on all 234 patches (2 dung pats x 2 time points = 936 dung pats in total). Cow dung was provided by Naturlandhof Peter in Schwebheim, Germany, an organic dairy farm, where cows were not treated with antibiotics or anthelmintics. Since dung was collected over several weeks in 2022 and 2023 and had to be processed immediately after collection, dung consistency varied slightly between collections.

Each collection was therefore provided with an ID (DungID), and it was taken care that for experiments conducted in each forest site, dung pats with the same DungID were used to ensure comparability between forest districts. After collection, the dung was portioned into dung pats of 400 – 420 g and stored frozen at – 20°C. Dung pats were defrosted 24 h before each exposure experiment.

Dung pats were placed 5 m apart from each other directly on the soil in the center of the patch. Both dung pats were covered with a small metal cage (16 x 30 x 40 cm) with an integrated rain cover to prevent access of vertebrates and to exclude the erosion of dung by rain (Figure S3). One of the cages was additionally wrapped in an insect net (mesh size 1 mm), preventing insect access both from above and below ground. Dung pats were weighed individually before exposure to patch conditions. After two days, the remaining dung pats were collected and dried for five days at 100°C. Dung removal was determined by calculating the relationship between the dry weight of the dung pats after exposure and the initial dry weight. The initial dry weight was estimated by calculating the dry weight/wet weight ratios of five to ten subsamples per dung collection and multiplying this ratio by the initial wet weight of the exposed dung samples. For each season, standard deviations of removal rates were calculated within each district to compare variances in removal rates and their relation to forest structure.

### Dung beetle assemblages

Dung beetles were collected directly after each dung exposure experiment by placing one pitfall trap baited with cow dung on each patch (Gebert et al., 2020, 2022). As baits, tea filter bags filled with 80 g of cow dung were used, which were stored frozen until 24 h before being used in the field. Baits were fastened to a wooden stick, which was secured in the ground next to the pitfall trap so that the bait was hanging freely above the opening of the trap. As traps, 400 ml plastic cups (volume: 473 cm^3^) were filled with a mixture of water and odorless dish soap. After 48 hours, specimens were removed from the trap and stored in 70 % ethanol for further processing. Permits were obtained from the respective regional authorities (Regierung von Niederbayern; Regierung von Unterfranken; Ministerium für Umwelt, Klima, Mobilität, Agrar und Verbraucherschutz Saarland; Landesamt für Umwelt Schleswig-Holstein). No further ethics approval was required. In the laboratory, each sample was searched for dung beetles of the families Geotrupidae and Scarabaeidae, which were identified to species level. Nomenclature follows the catalogue of Palaearctic Coleoptera (Löbl & Löbl, 2016).

To obtain the average biomass of each dung beetle species, five individuals from each species and each region were dried at 50°C for seven days, and their weights were averaged. If fewer than five individuals of a species were caught, all individuals were weighed.

### Environmental variables and defecation rates

For each sample, the average air temperature at ground level (measured at 15 cm above ground) of the sample month for each study patch was calculated from temperature data of four TOMST TMS-4 data loggers, which were placed on each patch. Five-year averages of cumulated precipitation were derived from yearly gridded datasets of the German Meteorological Service (Source: Deutscher Wetter Dienst DWD). Since temperate forest dung beetles have been shown to prefer sandy soils (von Hoermann et al., 2022) we included clay-to-sand ratios to control for differences in forest soils between study regions/forest districts. Clay-to-sand ratios were calculated from clay and sand percentages of soil core data from the first mineral soil layer of each forest patch collected in 2024 (Cesarz & Gulder, 2025).

Data on mammal species occurrences in 2022 and 2023 were extracted per forest district from camera traps positioned continuously from March to July on all study plots (Müller, 2025).

Mammal occurrences were converted into defecation rates by raising the biomass of each mammal species to the power of ¾, multiplying it by the frequency of each species on the camera traps in each district, respectively, and summing up the values across all species (Gebert et al., 2020). Mean defecation rates per patch were calculated by dividing forest district values by the number of study patches contained in one district.

## Statistical analysis

All statistical analysis was conducted with R 4.4.1 (R Core Team, 2024). The package tidyverse (Wickham et al., 2019) was used for data handling and visualization, and glmmTMB (Brooks et al., 2017) for modelling.

Clay-to-sand ratios and climate categories (warm climate/cold climate) were included in all models as control variables representing climatic and soil conditions.

We calculated dung beetle α-diversity as the number of species per sampling and study patch. For each forest district, we calculated γ-diversity, additive and multiplicative β-diversity. γ-diversity was calculated as the number of all dung beetle species in the district. For the calculation of γ-and β-diversity, mean α-diversity was calculated by pooling the species records from both sampling seasons and averaging them across the patches of the forest district. Additive and multiplicative partitioning of β-diversity provide complementary perspectives on species turnover among sites, allowing us to capture both the magnitude and the relative extent of species turnover among patches within each forest district. While additive β-diversity (additive β-diversity = γ-diversity – mean α-diversity) quantifies the absolute gain in species richness from patch to landscape scale, multiplicative β-diversity (multiplicative β-diversity = γ-diversity / mean α-diversity) describes the proportional increase in species richness relative to mean patch diversity.

For modelling the effect of forest structural complexity (forest district), separate generalized linear mixed effect models (GLMMs) were fitted for dung removal with insect access (no insect net), α-diversity (n = 468), β-diversity (n = 22), and γ-diversity (n = 22), and a random effect of forest site was added to each model. To test if forest structural complexity (forest district) increased differences in removal rates between forest patches, we additionally calculated standard deviations (SD) of removal rates with insect access at the forest district level for each season and district combination (n = 44).

Effects of canopy gaps on α-diversity and dung removal with insect access were tested by grouping study patches by their openness due to management into the categories “aggregated” (canopy gap), “distributed” (removed trees evenly distributed across the patch) and “control” (closed canopy) and fitting a GLMM testing the effect of “ESBC-treatment” for each response variable, respectively. Since there were control patches in each forest district, a random effect of forest district nested in forest site was included in each model.

For testing the effect of insect exclusion on removal rates (n=923) we fitted a GLMM including a random effect for study patch nested in forest site, since dung pats were exposed in pairs on each patch within the forest sites (Figure 1). A total of 13 samples were destroyed by wild boars and could not be included in the analysis.

We fitted GLMMs to model the effects of temperature during the sampling period and soil clay-to-sand ratios on dung removal rates (random factor DungID) and dung beetle biomass (n=468).

To investigate the role of dung beetles in dung removal we used separate GLMMs to model the effects of dung beetle α-diversity, abundance, and average biomass on dung removal rates with insect access, including DungID as a random factor in each model to account for differences in dung consistency across samples.

We further tested the effects of climate category and forest district on mean mammalian defecation rates and the relationships between mean defecation rates and mean α-diversity as well as mean abundance respectively, using linear models.

All models of dung removal rates were modelled using a beta distribution, while SDs of removal rates were modelled using a Gaussian distribution. Biomass was modelled using a zero-inflated log-normal distribution or a zero-inflated beta distribution, and beetle diversity and abundance data were modelled with a Poisson or negative binomial distribution, and in case of zero inflation with a truncated Poisson distribution.

For all the above-mentioned models, model coefficients were extracted using Anova type II to obtain main effects adjusted for other predictors when no interaction terms were included. For models including interaction terms, ANOVA Type III was used, which tests each main and interaction effect independently, without assuming hierarchical nesting. Contrasts for categorical variables with more than two levels, as well as for interactions, were extracted using Tukey’s HSD test for post – hoc analysis of group differences from the package emmeans (Lenth, 2024). All model diagnostics were performed using the package DHARMa (Hartig, 2024). Confidence intervals for estimated marginal effects were calculated with ggeffects (Lüdecke, 2018).

## Results

### Effects of structural complexity in forests on dung beetle diversity and dung removal

We found no significant differences between the structurally homogenous control districts and structurally heterogeneous ESBC districts for γ-, and additive β-diversity (Figure 2a, c, Table S1). However, multiplicative β-diversity and α-diversity were significantly lower in ESBC forest districts (Figure 2b, d, Table S1).

**Figure 2.**
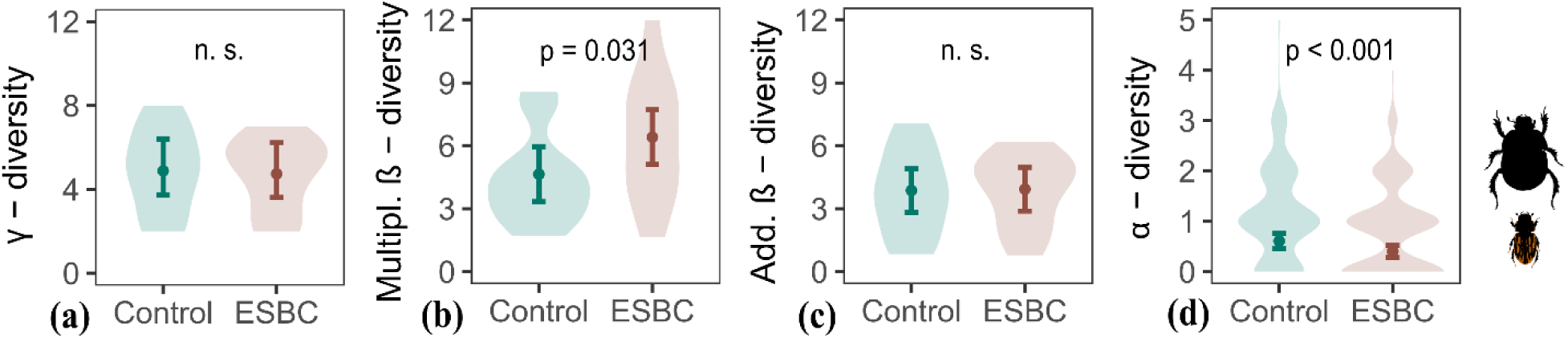
γ-diversity **(a)**, multiplicative β-diversity **(b)**, additive β-diversity **(c)**, and α-diversity **(d)** of dung beetle species in homogenous control districts (Control) and forest districts with enhanced structural complexity (ESBC). Error bars represent 95 % confidence intervals, and points represent estimates of marginal means. Violin plots show the raw data density distribution.

In cold climates, forest districts with a heterogeneous habitat structure at the landscape level (ESBC-districts) had significantly lower mean dung removal rates (with insect access) than control districts in both sampling seasons. In warm climates, no effect of enhanced structural complexity of forests on mean removal rates was observed (Figure 3a, Table S2, S3). The variation in dung removal rates (for samples with insect access) was not different between control and ESBC-districts, meaning that forest districts with increased structural complexity did not show a broader range of decomposition rates (Figure 3b, Table S2).

**Figure 3.**
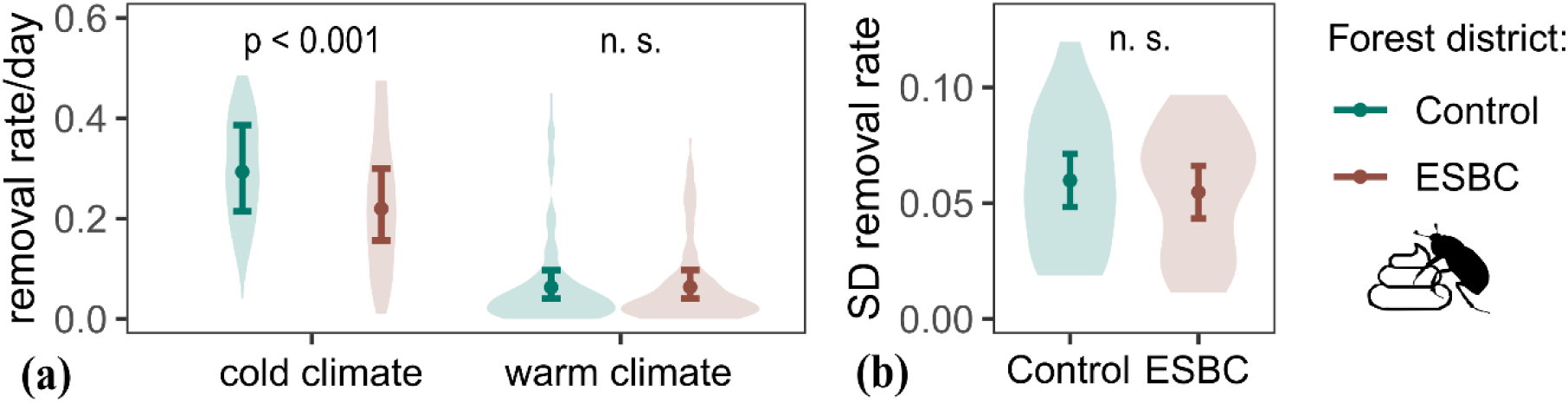
Dung removal rates **(a)** and standard deviation of dung removal rates per forest district **(b)** of dung pats with insect access in homogenous control forest districts (Control) and forest districts with enhanced structural complexity (ESBC). Error bars represent 95 % confidence intervals and points represent estimates of marginal means. Violin plots show density plots of the raw data distribution.

### Effect of canopy coverage on local dung beetle assemblages and dung removal

The reduced α-diversity and multiplicative β-diversity in ESBC districts suggested that dung beetle species and dung removal were reduced on some forest patches (Figures 2, 3). Indeed, α-diversity was reduced in forest patches where the forest canopy had been manipulated. This effect was especially strong in warm climates where α-diversity was strongly reduced in ESBC-treatments with continuous canopy gaps (aggregated treatment) (Figure 4a, Table S4, S5).

**Figure 4.**
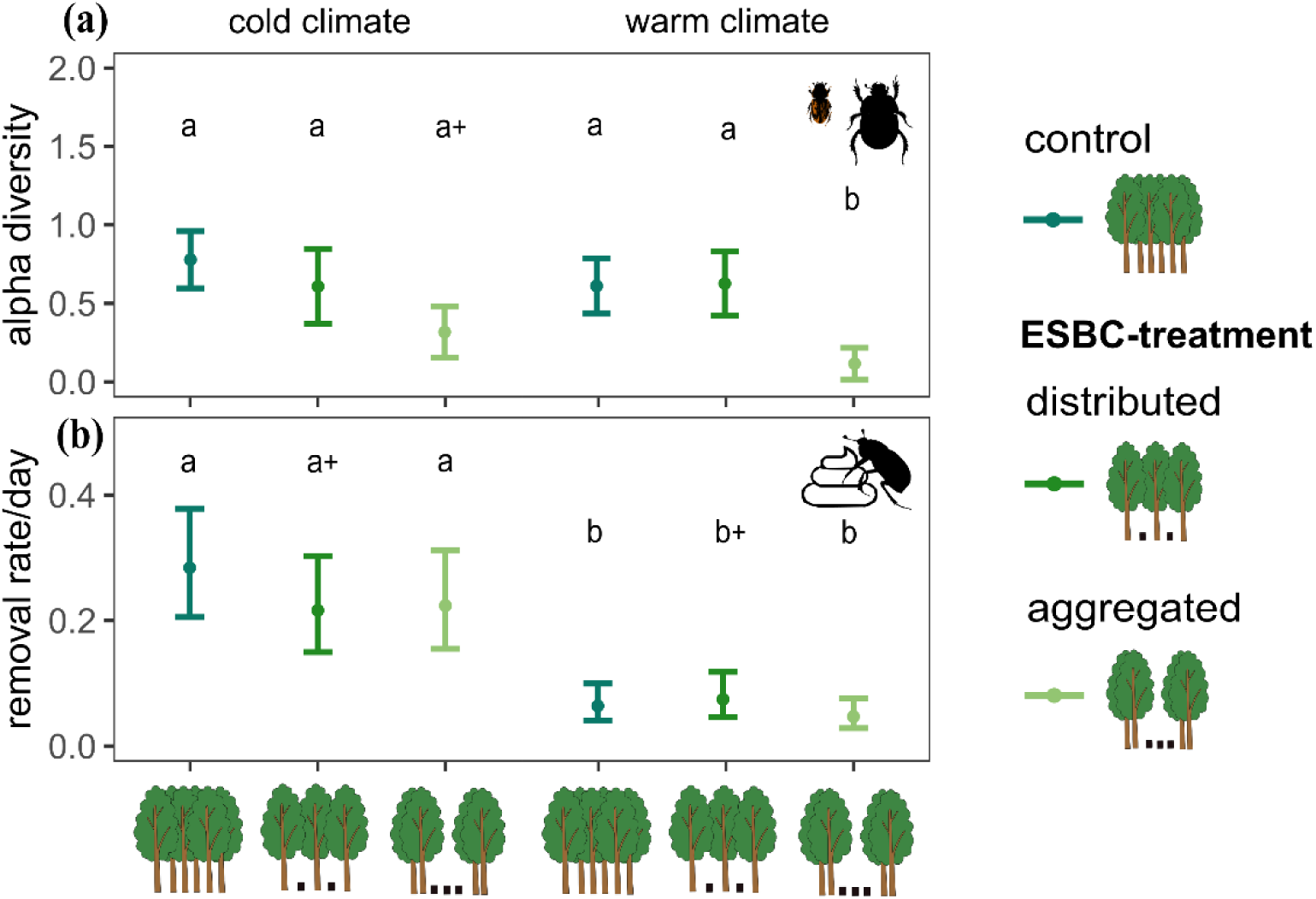
Dung beetle α-diversity **(a)** and removal rates of cow dung **(b)** in homogeneous control forest study patches (control) and ESBC-treatment forest patches with 30 % trees manipulated either distributed over the patch (distributed) or only in the patch center (aggregated). Letters indicate differences between groups, while + indicates marginally significant effect (0.05 < p < 0.1).

Removal rates (with insect access) did not show significant differences between the different treatments, but were strongly reduced in warmer climates in all treatments (Figure 4b, Table S4, S5).

### Contribution of insects and climate to dung removal

In colder climates, removal rates of dung pats that were covered by an insect net (insect exclusion) were reduced by 77 - 91 % compared with dung pats that were open to invertebrate detritivores (insect access) (Figure 5, Table S6, S7). In warmer climates, this effect occurred only in spring (67 % lower with insect exclusion), while during late summer, removal rates of dung pats with and without insect showed no difference (Figure 5, Table S7).

**Figure 5.**
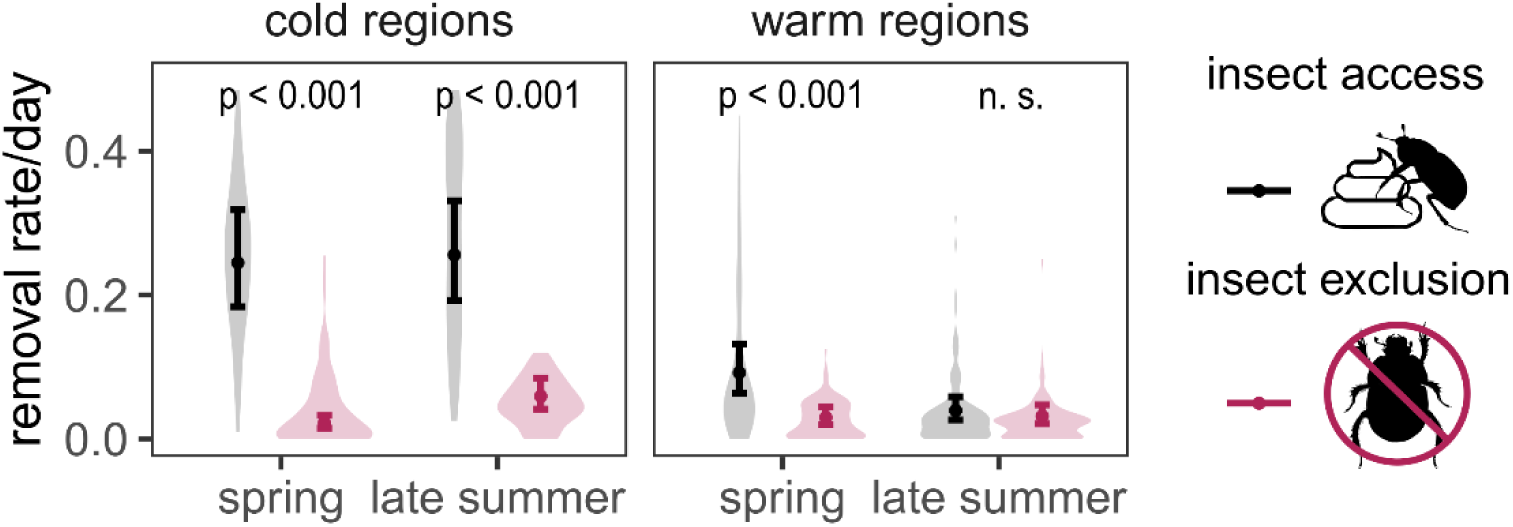
Dung removal rates, measured as the fraction of dung dry weight removed per day for dung pats with and without insect access for forest sites situated in warm and cold climate categories. Error bars represent 95 % confidence intervals, and points represent estimates of marginal means. Violin plots show the raw data distribution.

Dung removal rates of dung pats with insect access showed a strong negative relationship with mean air temperature (Figure S4, Table S8). This was not the case for dung pats under insect exclusion, where we found no relationship between mean air temperatures and removal rates, suggesting that the temperature dependence of dung removal was driven by a higher dung beetle activity in cold climates (Figure S4, Table S8).

We recorded 2730 dung beetle individuals from 12 species (Table S9). All recorded species were forest specialists or habitat generalists (Frank et al., 2017; Staab et al., 2022). Average dung beetle biomass per study patch decreased with increasing temperature and increasing proportions of sand in the soil (Figure 6a, Table S10). In cold climates, dung beetle abundance and biomass were dominated by the large tunneller species *Anoplotrupes stercorosus* (Geotrupidae), while small dweller species, comprised most dung beetle individuals in warm climates (Figure 6b).

**Figure 6.**
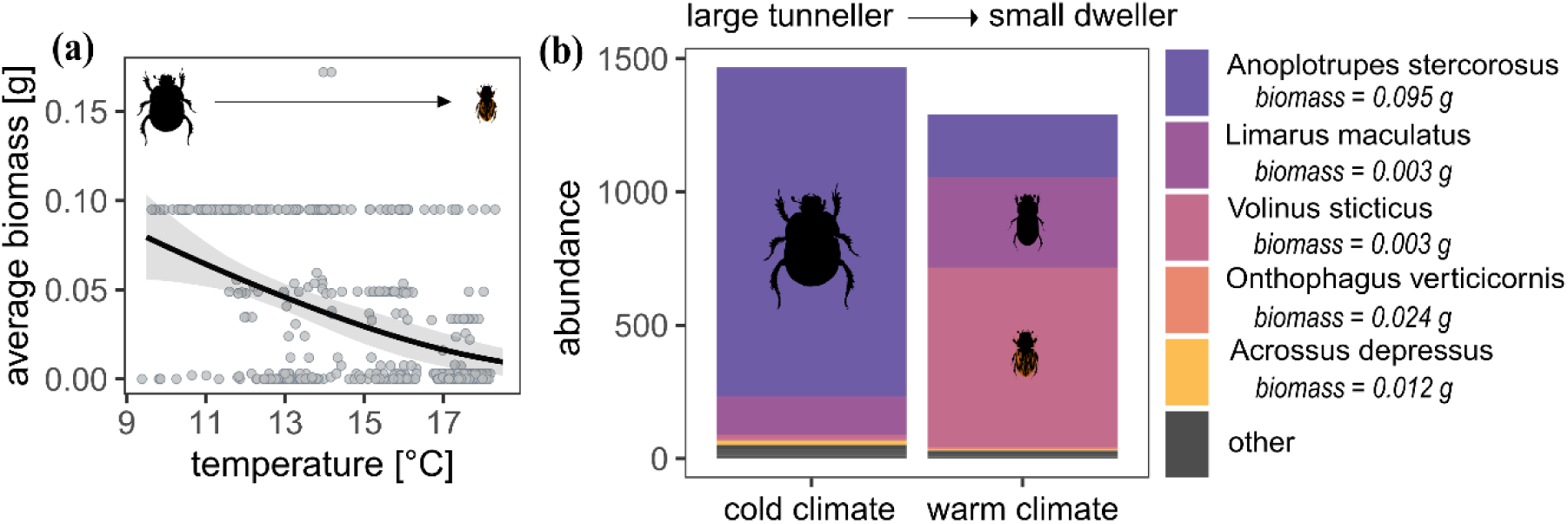
Effects of temperature on average dung beetle biomass per patch **(a)** and abundance **(b)** in study regions separated in cold and warm climate categories. Grey areas represent 95% confidence intervals, and lines represent estimates of marginal effects. Points show raw data (a).

Tunneller bury brood balls underneath the dung pats, while dwellers brood inside the dung. Dung removal rates were positively related to average beetle biomass, but not to α-diversity or abundance of dung beetles (Figure S5, Table S11).

### Effects of mammalian activity on dung beetle communities

Mammalian defecation rates did not differ between control forest districts and ESBC forest districts and did not explain variations in dung beetle abundance or α-diversity. Additionally, mean defecation rates were higher in regions with warmer climates (Figure 7a-d, Table S12), suggesting that it is not the amount of feces of mammals that drives dung removal by dung beetles or that the amount is not a limiting factor.

**Figure 7.**
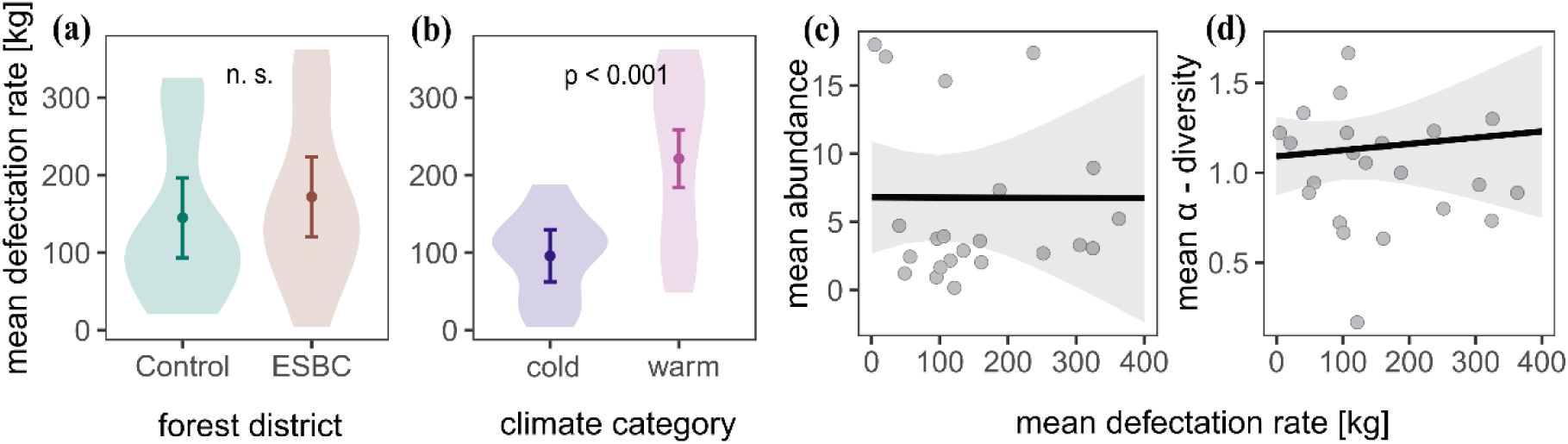
Effects of forest structure **(a)** and climate **(b)** on mean defecation rates of mammals in control and ESBC-districts. Relationship of mean mammal defecation with mean abundance **(c)** and mean α-diversity of dung beetles **(d)** for each forest district. Grey areas represent 95% confidence intervals, and lines represent estimates of marginal effects. Points show raw data, and violin plots show density distributions of raw data.

## Discussion

The enhancement of structural heterogeneity in forests (ESBC) did neither foster higher dung removal rates nor higher β-and γ-diversity of dung beetles. On the contrary, canopy gaps as part of the ESBC forest management in interaction with higher air temperatures negatively affected dung beetle α-diversity and abundance. Dung removal was driven by average dung beetle biomass, which decreased with increasing temperature due to a species turnover from large tunneller dung beetles to small dweller species. Additionally, dung beetle diversity and abundance were not controlled by the availability of mammalian dung but by climate and forest structure.

### Effects of structural complexity in forests on dung beetle diversity and dung removal

Due to complementarity in the climate and habitat niches of dung beetles, we expected that a higher dissimilarity in forest structure between patches, as created by the ESBC-management, would lead to a higher β-and γ-diversity at the landscape scale. Contrary to our expectations, dung beetles did not profit in terms of α-, β-, or γ-diversity from increased structural heterogeneity in our ESBC forest landscapes. Multiplicative β-diversity was higher in ESBC districts, however, this was caused by a decline in α-diversity in patches with an open canopy, not by species turnover. Decreases in forest dung beetle species in canopy gaps were not compensated by influx of open-habitat specialists.

The large-scale forest manipulation experiment used in this study allowed to assess the effect of increased heterogeneity in the forest landscape on temperate dung beetle communities at α-, β-, and γ-scales for the first time. Our findings suggest that, while other groups might respond positively to increased structural heterogeneity in production forests (Doerfler et al., 2018; Dove & Keeton, 2015; Lettenmaier et al., 2022; Rothacher et al., 2025), temperate dung beetle communities are adapted to relatively cool, closed canopy forests and do not profit from management strategies such as ESBC at the β-and γ-level.

Moreover, we found negative effects of larger canopy openings on dung beetle α-diversity in line with previous studies in both temperate and tropical forests that showed negative impacts of intensive logging regimes and clear-cutting on dung beetle communities (Frank et al., 2017; Horgan, 2005; Sánchez-de-Jesús et al., 2016; Slade et al., 2011, Staab et al. 2022). In our study, negative effects of canopy openness on dung beetle α-diversity were strongest in warmer climates, showing that forest dung beetles do forage in more open forest habitats, when climatic conditions are suitable, but avoid accessing these habitats under increasing temperatures. Gap effects might therefore be driven by temperature, both by increasing the physiological stress of dung beetles through increased water loss and by decreasing dung resource attractiveness through desiccation (Holter, 2016; Staab et al., 2022). Even though we found a community turnover from large tunneller species to small dweller species along the climate gradient, combinations of high temperatures and canopy gaps influenced both functional groups alike.

Thus, the lowered dung beetle activity in open canopy forests can be related to special climate and habitat niche characteristics of the dung beetle community dominating today’s Central European forests.

We did not find an additional influx of dung beetle species specialized on more open habitats to forest patches dominated by canopy gaps (aggregated treatment). Instead, dung beetle assemblages in these open forests were a subset of the dung beetle communities found in closed forests. It is possible that the experimental canopy openings were not large enough to attract open habitat specialists like *Geotrupes spiniger* or *G. mutator,* the largest tunneller species on grassland pastures. However, over the past decades, open habitat specialists have declined due to agricultural intensification, which has led to a decline in grazing livestock and a loss of habitat. Studies have found lower densities of large tunneller species in temperate grasslands as compared to forests, which could explain, why large grassland species were not recruited to the canopy gap (Buse & Entling, 2020; Frank et al., 2017).

Under future climate change, an increased frequency of forest disturbances leading to more open forests in combination with rising temperatures might put the current temperate forest dung beetle community at risk, forcing species to retreat to higher latitudes and altitudes (Bednar-Friedl et al., 2022). It is therefore crucial that, in the effort of conserving biodiversity in production forests through targeted management, we also focus on preserving forest stands with dense forest canopies in combination with more open areas that might benefit other groups.

### Contribution of insects and climate to dung removal

We found that, on average, insects were responsible for 86% of the dung removal in Central European production forests. However, dung removal through insects was negatively related to temperature. Under high temperatures (in warm climates and in late summer), the contribution of insects to dung removal was no longer significant, revealing the significant but climate-dependent contribution of insects to dung removal in managed forest ecosystems. This pattern could be explained by a species turnover along the climate gradient from communities dominated by the large tunneller dung beetle species *Anoplotrupes stercorosus* to communities mostly comprised of small dweller dung beetle species. Dung removal was therefore driven by the presence of *A. stercorosus* and was correlated with average dung beetle biomass per forest patch, but not with α-diversity and abundance.

*A. stercorosus* is a large forest-adapted beetle of the family Geotrupidae, which buries brood balls made of dung in tunnel systems near the dung pat and is known for its high dung removal efficiency and abundance in temperate forests (Frank et al., 2017; Nervo et al., 2014). It not only plays an essential role in dung removal but also supports bioturbation and soil aeration by moving large quantities of the forest soil (Nichols et al., 2008).

We expected that dung removal would be higher in habitats with higher temperatures, since both field and laboratory experiments showed increased dung removal through dung beetles, and *A. stercorosus* specifically, under warmer temperatures (Nervo et al., 2024; von Hoermann et al., 2022). Additionally, we expected dung removal to increase on sandy soils, as loose soil facilitates tunnelling behavior (von Hoermann et al. 2022). Consistent with earlier findings, we found positive effects of soils with high proportions of sand on dung beetle biomass. Against expectations, our findings suggest that increased temperatures negatively affect dung removal and the abundance of *A. stercorosus* when assessing dung beetle communities over large scales in field conditions. Nervo et al. (2024) tested dung removal from different large tunneller species by varying temperatures according to climate change scenarios but keeping all other environmental variables constant. Under field conditions, this is, however, rarely the case. In temperate regions, temperature and air moisture are usually negatively related, and possibly, a combination of climatic factors created conditions unfavorable for *A. stercorosus*, leading to reduced abundances of this species under warmer conditions. Larger dung beetles are more vulnerable to water loss, due to larger-sized spiracles, which could explain the species turnover we observed along the climate gradient (Nervo et al., 2021).

Our findings further solidify previous suggestions of a species identity effect of *A. stercorosus* on dung removal in temperate forest, showing that patterns are consistent across seasons and along a climate gradient (Staab et al, 2022). Because of the lack of many other abundant and functionally redundant tunneller species, temperate forest ecosystems might be especially vulnerable to the loss of this one dominant species (Frank et al., 2017). With droughts and extreme temperatures projected to increase in the future, it is likely that *A. stercorosus* will suffer increased environmental pressures, and its distributional range in Central Europe will decrease (Bednar-Friedl et al., 2022). Thermophilic dung beetle species from the Mediterranean Region are predicted to shift their ranges northwards in the next decades, however, whether species achieve the shift in latitude depends on the availability of suitable habitat corridors, which are decreasing under continued land use change (Dortel et al., 2013).

### Effects of mammalian activity on dung beetle communities

Against expectations, dung beetle richness and abundance were not correlated with mammalian defecation rates. In contrast to dung removal rates, which decreased with increasing temperature, defecation rates were higher in warmer climates. Since negative effects of mammal declines due to persistent game hunting on dung beetle communities have been shown by several studies, we expected that dung beetle diversity and abundance would increase with increased dung resource availability (Buse et al., 2021; Nichols et al., 2009; Raine & Slade, 2019). However, in German production forests, populations of large mammals such as roe deer and wild boars are not declining, thus dung availability may not limiting in these ecosystems (Greiser et al., 2023). Our findings suggest that if dung beetle communities are not resource-limited, abiotic factors such as climate and soil texture are more important for constraining dung beetle abundance and diversity. Similar results were reported by Gerbert et al. (2020) along an elevational gradient at Mt.

Kilimanjaro, where temperature rather than dung resources predicted dung beetle diversity.

## Conclusion

The current dung beetle community in Central European production forests does not profit from a forest management that enhances structural complexity at the landscape scale. Temperate forest dung beetles have a restricted climatic niche and will therefore likely suffer negative consequences from increasing temperatures in the future. Changes in forest structure that lead to an opening of the forest canopy might exacerbate these negative effects from climate change.

Since forests face increased frequencies of natural disturbances that create large canopy openings, it is crucial for forest management to preserve closed-canopy forest stands, which would buffer temperature extremes and protect temperate dung beetle communities, and maintain the ecosystem functions they provide. Further research into the role of *A. stercorosus* in temperate forest ecosystems is needed to assess the impact of local extinctions on ecosystem functioning. Additionally, future studies should consider monitoring range shifts of thermophilic dung beetles and assessing their potential for filling the functional gap left by retreating temperate tunneller species.

## Author contributions

Marcell K. Peters, Michael Scherer-Lorenzen, Jörg Müller and Johanna Asch conceived the ideas and designed methodology; Johanna Asch, Julia Rothacher, Clara Wild, Kerstin Pierick, Orsi Decker, Nico Daume and Simone Cesarz collected the data; Jörn Buse advised on dung beetle identification; Johanna Asch analysed the data; Johanna Asch, Marcell K. Peters and Michael Scherer-Lorenzen interpreted the results; Johanna Asch wrote the original draft. All authors contributed critically to the drafts and gave final approval for publication.

## Data availability statement

The data is provided as supplementary files for reviewers and will be uploaded to a publicly available database (Zenodo) upon acceptance.

## Acknowledgements

We thank Anna-Lena Schleicher, Louis Puille, Sara Plesser, Franziska Asch, Leonie Feik, Pia Wapler, Lea Schamberger, Maria Camilla de la Hoz, Matein Weber, Matthias Weid, Carolin Moser, Lucas Gscheidle, Jean-Leonard Stör, Rabea Klümpers, Jens Schlüter and the team at the Bavarian Forest National Park for assistance in the field and laboratory, as well as Michael Junginger and Sonja Kümmet for their assistance with field work coordination and data management. We further thank Wolfgang Peter and his family from Naturlandhof Peter in Schwebheim for providing us with cow dung. Funding for the BETA-FOR research group was provided by the Deutsche Forschungsgemeinschaft (DFG, German Research Foundation, FOR 5375) – 459717468, with additional support from the Julius-Maximilians-Universität Würzburg (JMU).

