## supplementary material for "Dung beetles do not profit from enhanced spatial heterogeneity in production forests: a large-scale forest manipulation experiment"

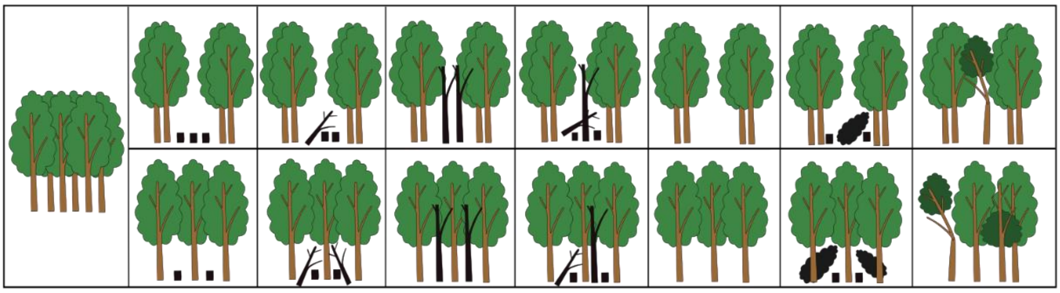

**Figure S1.** Graphical representation of forest structure manipulations of ESBC forest district in the study region University Forest. Black color represents different dead wood manipulations (for a detailed explanation of different dead wood manipulations see Müller et al. (2022).

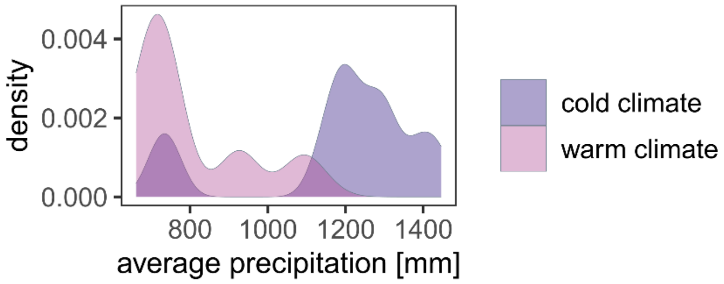


**Figure S2.** Density distributions of 5-year average precipitation sums in mm on 234 study plots separated into cold and warm climate regions.

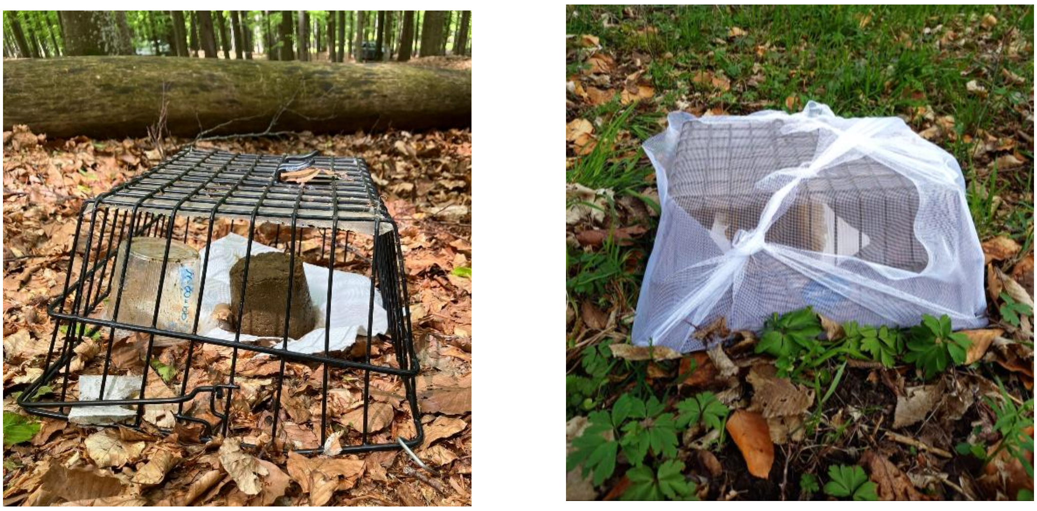

**(a)** **(b)**

**Figure S3.** Example of dung samples placed on each forest patch either in an open metal cage to allow insect access (a) or covered by an insect net (1mm mesh size) to exclude insects (b).

**Table S1.** Effects of enhanced structural complexity (ESBC) in forests (forest district) on gamma, additive beta, multiplicative beta modelled with separate generalized linear mixed-effects model (GLMMs) and alpha diversity modelled with a zero-inflated GLMM (both model components are reported). Each model included climate region and clay-to-sand ratios as control variables and modelled alpha diversity additionally included season as a control.

| Model | Variable | Chi-sq | Df | p.value |
|---|---|---|---|---|
| gamma diversity | **Forest district** | 0.023 | 1 | 0.023 |
| | Climate region | 1.025 | 1 | 0.311 |
| | Clay_Sand | 0.218 | 1 | 0.641 |
| additive beta diversity | **Forest district** | 0.053 | 1 | 0.817 |
| | Climate region | 3.934 | 1 | 0.047 |
| | Clay_Sand | 0.860 | 1 | 0.354 |
| multiplicative beta diversity | **Forest district** | 4.660 | 1 | 0.031 |
| | Climate region | 9.071 | 1 | 0.003 |
| | Clay_Sand | 3.429 | 1 | 0.064 |
| alpha diversity *conditional component* | **Forest district** | 1.455 | 1 | 0.228 |
| | Climate region | 0.606 | 1 | 0.436 |
| | Season | 29.393 | 1 | <0.001 |
| | Clay_Sand | 2.358 | 1 | 0.125 |
| alpha diversity *zero-inflated component* | **Forest district** | 12.885 | 1 | <0.001 |
| | Climate region | 7.448 | 1 | 0.006 |
| | Season | 0.098 | 1 | 0.754 |
| | Clay_Sand | 0.448 | 1 | 0.503 |

**Table S2.** Effects of ESBC-management (forest district) and climate on dung removal rates and standard deviations of dung removal rates modelled with generalized linear mixed-effects model. Each model included climate region, season and clay-to-sand ratios as control variables.

| Model | Variable | Chi-sq | Df | p.value |
|---|---|---|---|---|
| Dung removal rates | **Forest district** | 23.587 | 1 | **<0.001** |
| | Climate region | 31.849 | 1 | <0.001 |
| | season | 16.676 | 1 | <0.001 |
| | Clay_Sand | 0.038 | 1 | 0.846 |
| | **Forest district: Climate region** | 8.742 | 1 | **0.003** |
| Standard deviations of dung removal rates | **Forest district** | 0.974 | 1 | **0.324** |
| | season | 3.498 | 1 | 0.061 |
| | Climate region | 1.654 | 1 | 0.198 |
| | Clay_Sand | 4.639 | 1 | 0.031 |

**Table S3.** Pairwise contrasts of interactive effects of climate region and forest district on dung removal rates produced with a Tukey's HSD post-hoc test (extends Table S8).

| Control District vs. ESBC District | odds.ratio | SE | z.ratio | p.value |
|---|---|---|---|---|
| Cold regions | 1.491 | 0.102 | 5.86 | <0.001 |
| Warm regions | 1.028 | 0.099 | 0.28 | 0.992 |

**Table S4.** Results of separate general mixed effect models (GLMMS) for the effects of experimental manipulations of the forest canopy (treatment_canopy) in interaction with climate on alpha diversity and dung removal rates. Alpha diversity was modelled with a zero-inflated GLMM, both model components are reported. Each model included season and clay-to-sand ratios as control variables.

| Model | Variable | Chi-sq | Df | p.value |
|---|---|---|---|---|
| Alpha diversity *Conditional component* | treatment_canopy | 3.155 | 2 | 0.206 |
| | climate region | 0.531 | 1 | 0.466 |
| | season | 28.786 | 1 | <0.001 |
| | Clay_Sand | 2.935 | 1 | 0.087 |
| | treatment_canopy:climate region | 0.324 | 2 | 0.851 |
| Alpha diversity *Zero-inflated component* | treatment_canopy | 5.626 | 2 | 0.06 |
| | climate region | 11.059 | 1 | 0.001 |
| | season | 0.109 | 1 | 0.741 |
| | Clay_Sand | 0.011 | 1 | 0.918 |
| | treatment_canopy:climate region | 11.956 | 2 | 0.003 |
| Removal rates | treatment_canopy | 7.672 | 2 | 0.022 |
| | climate region | 28.273 | 1 | <0.001 |
| | season | 16.747 | 1 | <0.001 |
| | Clay_Sand | 0.040 | 1 | 0.841 |
| | treatment_canopy:climate region | 9.406 | 2 | 0.009 |

**Table S5.** Pairwise contrasts of interactive effects of climate region and experimental manipulations of the forest canopy on dung beetle alpha diversity and dung removal rates produced with a Tukey's HSD post-hoc test. Closed canopy: control, even removal of trees: distributed, clustered removal of trees i.e. canopy gap: aggregated. Extends Table S10.

| Model | Climatic region | contrasts | estimate | SE | z.ratio | p.value |
|---|---|---|---|---|---|---|
| Alpha diversity | Cold regions | control vs. distributed | 0.279 | 0.217 | 1.285 | 0.794 |
| | | control vs. aggregated | 0.460 | 0.172 | 2.670 | 0.081 |
| | | distributed vs. aggregated | 0.181 | 0.184 | 0.987 | 0.922 |
| | Warm regions | control vs. distributed | -0.158 | 0.205 | -0.771 | 0.972 |
| | | control vs. aggregated | 0.622 | 0.195 | 3.181 | 0.018 |
| | | distributed vs. aggregated | 0.779 | 0.148 | 5.279 | <0.001 |
| Dung removal | Cold regions | control vs. distributed | 1.448 | 0.202 | 2.657 | 0.084 |
| | | control vs. aggregated | 1.377 | 0.191 | 2.300 | 0.194 |
| | | distributed vs. aggregated | 0.951 | 0.112 | -0.425 | 0.998 |
| | Warm regions | control vs. distributed | 0.849 | 0.153 | -0.907 | 0.945 |
| | | control vs. aggregated | 1.387 | 0.256 | 1.772 | 0.484 |
| | | distributed vs. aggregated | 1.634 | 0.254 | 3.161 | 0.02 |

**Table S6.** Results of a general mixed effect model for the effects of insect exclusion, climate region, season and clay-to-sand ratios on dung removal rates.

| Variable | Chi-sq | Df | p.value |
|---|---:|---:|---|
| Insect exclusion | 691.896 | 1 | <0.001 |
| Climate region | 17.504 | 1 | <0.001 |
| Season | 0.703 | 1 | 0.402 |
| Clay_Sand | 0.797 | 1 | 0.372 |
| Insect exclusion:climate region | 189.203 | 1 | <0.001 |
| Climate region:season | 86.715 | 1 | <0.001 |
| Insect exclusion:season | 80.186 | 1 | <0.001 |

**Table S7.** Pairwise contrasts of interactive effects of insect exclusion, climate region and season on dung removal rates produced with a Tukey's HSD post-hoc test (extends Table S1).

| Insect access vs. exclusion | season | odds.ratio | SE | z.ratio | p.value |
|---|---|---:|---:|---:|---:|
| Cold regions (climate region) | spring | 14.357 | 1.454 | 26.304 | <0.001 |
| | late summer | 5.450 | 0.470 | 19.672 | <0.001 |
| Warm regions (climate region) | spring | 3.310 | 0.323 | 12.259 | <0.001 |
| | late summer | 1.256 | 0.121 | 2.373 | 0.254 |

**Table S8.** Results of a general mixed effect models for the effects of mean air temperature on dung removal rates with insect access (a) and insect exclusion (b). Each model included season and clay-to-sand ratios as control variables.

| Model | Variable | Chi-sq | Df | p.value |
|---|---|---|---|---|
| (a) dung removal insect access | mean.T | 110.403 | 1 | <0.001 |
| | season | 45.529 | 1 | <0.001 |
| | Clay_Sand | 26.602 | 1 | <0.001 |
| (b) dung removal insect exclusion | mean.T | 0.376 | 1 | 0.54 |
| | season | 11.423 | 1 | 0.001 |
| | Clay_Sand | 2.520 | 1 | 0.112 |

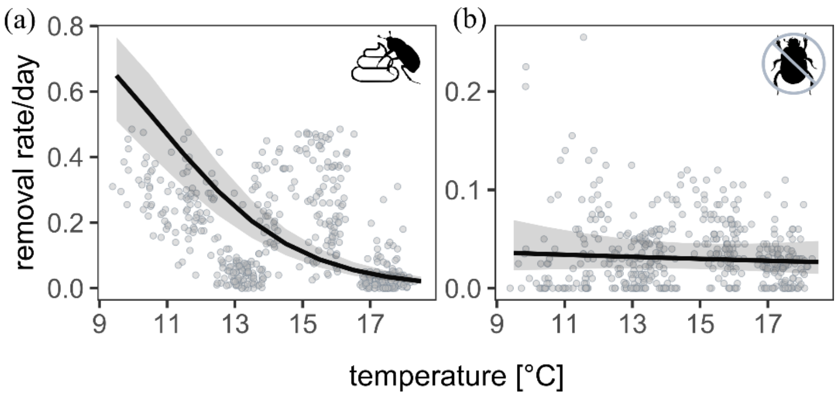


**Figure S4.** Dung removal rates as fraction removed per day of dung pats with insect access **(a)** and with insect exclusion **(b)** and their relationship with mean air temperature in °C. Grey areas represent 95% confidence intervals and lines represent estimates of marginal effects. Points show raw data.

**Table S9.** Dung beetle species collected in each study region and their percentage of the total abundance and biomass in each region.

| Region | Species | Abundance (%) | Biomass (%) |
|---|---|---|---|
| Bavarian Forest | *Acrossus depressus* | 1.17 | 0.17 |
| | *Acrossus rufipes* | 0.16 | 0.03 |
| | *Anoplotrupes stercorosus* | 86.78 | 99.38 |
| | *Calamosternus granarius* | 0.31 | 0.01 |
| | *Limarus maculatus* | 10.89 | 0.39 |
| | *Parammoecius corvinus* | 0.23 | <0.01 |
| | *Volinus sticticus* | 0.47 | 0.02 |
| Hunsrueck | *Acrossus depressus* | 2.17 | 0.44 |
| | *Anoplotrupes stercorosus* | 61.59 | 98.07 |
| | *Aphodius fimetarius (L. 1758)* | 2.90 | 0.39 |
| | *Calamosternus granarius* | 7.97 | 0.27 |
| | *Limarus maculatus* | 1.45 | 0.07 |
| | *Parammoecius corvinus* | 13.04 | 0.22 |
| | *Volinus sticticus* | 10.87 | 0.55 |
| Luebeck | *Anoplotrupes stercorosus* | 78.57 | 99.40 |
| | *Calamosternus granarius* | 14.29 | 0.38 |
| | *Limarus maculatus* | 4.76 | 0.19 |
| | *Parammoecius corvinus* | 2.38 | 0.03 |
| Passau | *Acrossus depressus* | 10.26 | 1.81 |
| | *Acrossus rufipes* | 2.56 | 0.60 |
| | *Anoplotrupes stercorosus* | 69.23 | 96.87 |
| | *Calamosternus granarius* | 2.56 | 0.08 |
| | *Limarus maculatus* | 5.13 | 0.23 |
| | *Onthophagus joannae* | 2.56 | 0.23 |
| | *Parammoecius corvinus* | 5.13 | 0.08 |
| | *Volinus sticticus* | 2.56 | 0.11 |
| Saarland | *Anoplotrupes stercorosus* | 93.55 | 95.57 |
| | *Aphodius fimetarius (L. 1758)* | 0.81 | 0.07 |
| | *Parammoecius corvinus* | 4.03 | 0.04 |
| | *Trypocopris vernalis* | 1.61 | 4.32 |
| University Forest | *Acrossus depressus* | 0.09 | 0.10 |
| | *Acrossus rufipes* | 0.27 | 0.40 |
| | *Anoplotrupes stercorosus* | 8.07 | 72.50 |
| | *Calamosternus granarius* | 0.09 | 0.02 |
| | *Limarus maculatus* | 30.08 | 8.53 |
| | *Limarus zenkeri* | 0.09 | 0.03 |
| | *Onthophagus verticicornis* | 0.62 | 1.41 |
| | *Parammoecius corvinus* | 0.98 | 0.09 |
| | *Volinus sticticus* | 59.72 | 16.93 |

**Table S10.** Results of both model components of a zero-inflated general mixed effect model for the effects of mean air temperature, clay-to-soil ratio and season on average dung beetle biomass.

| Component | Variable | Chi-sq | Df | p.value |
|---|---|---|---|---|
| conditional | mean.T | 8.518 | 1 | 0.004 |
| | season | 5.537 | 1 | 0.019 |
| | Clay_Sand | 28.921 | 1 | <0.001 |
| zero-inflated | mean.monthT | 14.97 | 1 | <0.001 |
| | season | 12.210 | 1 | <0.001 |
| | Clay_Sand | 0.164 | 1 | 0.686 |

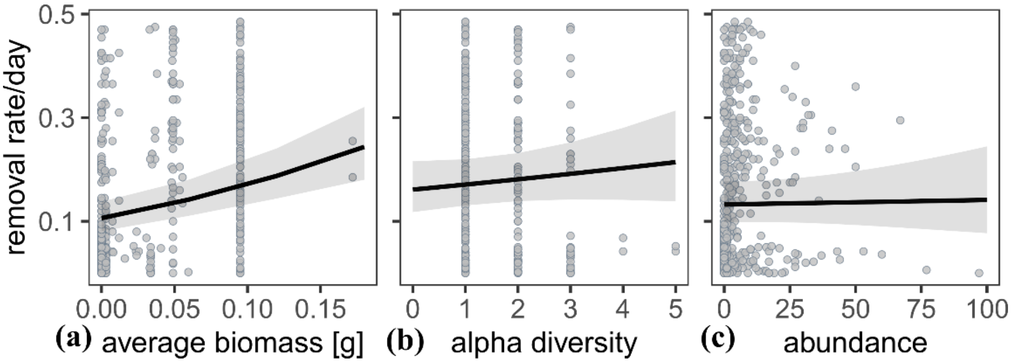


**Figure S5.** Relationships of dung beetles average biomass **(a),** alpha diversity **(b)** and abundance **(c)** with dung removal rates per day from dung beetles sampled with baited pitfall trap and dung pats placed on 234 study patches. Grey areas represent 95% confidence intervals and lines represent estimates of marginal effects. Points show raw data.

**Table S11.** Results of separate general mixed effect models for the effects of dung beetle average biomass **(a)**, alpha diversity **(b)**, and abundance **(c)** on dung removal rates. Each model included season, climate region and clay-to-sand ratios as control variables.

| model | Variable | Chi-sq | DF | p.value |
|---|---|---|---|---|
| (a) | **average biomass** | 32.892 | 1 | <0.001 |
| | Clay_Sand | 24.246 | 1 | <0.001 |
| | season | 50.666 | 1 | <0.001 |
| | mean.monthT | 106.056 | 1 | <0.001 |
| (b) | **alpha diversity** | 0.800 | 1 | 0.371 |
| | Clay_Sand | 29.845 | 1 | <0.001 |
| | season | 22.735 | 1 | <0.001 |
| | mean.monthT | 66.636 | 1 | <0.001 |
| (c) | **abundance** | 0.051 | 1 | 0.821 |
| | Clay_Sand | 26.560 | 1 | <0.001 |
| | season | 45.491 | 1 | <0.001 |
| | mean.monthT | 110.132 | 1 | <0.001 |

**Table S12.** Effects of experimental manipulations of the forest canopy (forest district) and climate region on mean defecation rates of mammals. Effects of mean defecation rates of mammals on mean dung beetle abundance and mean dung beetle alpha diversity. Effects modelled with generalized linear mixed-effects models.

| Model | Variable | Df | Chi-sq | p.value |
|---|---|---|---|---|
| Mean defecation | Forest district | 1 | 1.1169 | 0.29 |
| | Climate region | 1 | 9.09 | 0.003 |
| Mean abundance | Mean defecation | 1 | 0.0001 | 0.99 |
| Mean alpha diversity | Mean defecation | 1 | 0.216 | 0.642 |
